# Visual Image Reconstructed Without Semantics from Human Brain Activity Using Linear Image Decoders and Nonlinear Noise Suppression

**DOI:** 10.1101/2023.09.23.559147

**Authors:** Qiang Li

## Abstract

In recent years, substantial strides have been made in the field of visual image reconstruction, particularly in its capacity to generate high-quality visual representations from human brain activity while considering semantic information. This advancement not only enables the recreation of visual content but also provides valuable insights into the intricate processes occurring within high-order functional brain regions, contributing to a deeper understanding of brain function. However, considering fusion semantics in reconstructing visual images from brain activity involves semantic-to-image guide reconstruction and may ignore underlying neural computational mechanisms, which does not represent true reconstruction from brain activity. In response to this limitation, our study introduces a novel approach that combines linear mapping with nonlinear noise suppression to reconstruct visual images perceived by subjects based on their brain activity patterns. The primary challenge associated with linear mapping lies in its susceptibility to noise interference. To address this issue, we leverage a flexible denoised deep convolutional neural network, which can suppress noise from linear mapping. Our investigation encompasses linear mapping as well as the training of shallow and deep autoencoder denoised neural networks, including a pre-trained, state-of-the-art denoised neural network. The outcome of our study reveals that combining linear image decoding with nonlinear noise reduction significantly enhances the quality of reconstructed images from human brain activity. This suggests that our methodology holds promise for decoding intricate perceptual experiences directly from brain activity patterns without semantic information. Moreover, the model has strong neural explanatory power because it shares structural and functional similarities with the visual brain.

## I. Introduction

The human brain, a remarkably intricate organ, serves as the foundation for cognitive information processing, managing the diverse range of our daily experiences [1–3]. Visual perception is a key part of cognitive function and is closely connected to both Short-Term Memory^1^ and Long-Term Memory^2^. Short-term memory holds a limited amount of information briefly, while long-term memory retains knowledge over extended periods [4]. To elucidate the complex relationship between visual experiences and brain states, neural encoding and decoding models were employed to explain the statistical dependence based on features derived from both visual experience and neural signals [5].

In the early stages of this exploration, linear mapping decoders were a prominent choice [6]. However, the primary challenge associated with linear decoders lies in their vulnerability to noise, which can come from various sources. Firstly, noise can originate from intrinsic brain activity itself. The Blood-Oxygen-Level-Dependent (BOLD) signals, commonly employed in neuroimaging studies, are susceptible to both physiological noise stemming from bodily functions like heart rate variations [7], and physical noise introduced by the MRI machinery [8]. Secondly, linear decoders often have difficulty handling high-dimensional and nonlinear functional magnetic resonance imaging (fMRI) data [9, 10]. The complex nature of neural activity patterns requires more advanced decoding methods. Furthermore, the problem of noise is made worse by the natural redundancy in images. While this redundancy helps humans recognize images more easily, it creates a significant challenge for linear decoders, leading to noticeable noise in the reconstructed images.

With the advent of deep learning models, a transformative shift occurred, leading to a diverse array of nonlinear decoding models. This paradigm shift resulted in remarkable outcomes, including superior performance in reconstructing visual content from neural activity [11–20]. These advancements have collectively enhanced the understanding and application of nonlinear decoding models for reconstructing visual information from brain activity. Even though the nonlinear deep net models have nice performance in reconstructing visual images, they lack neural explanations. Most notably, the reconstructed images are often derived from brain activity patterns that do not perfectly align with the visual stimuli images. Furthermore, the presence of widespread noise, coming from various sources, represents a significant challenge. As we mentioned above, these noise sources encompass artifacts from scanning machines [8], intrinsic neural noise [21, 22], and even physiological factors like heart rate fluctuations [7]. Overcoming these challenges is a key focus in improving the accuracy of reconstructing visual images from brain activity. It is also important to recognize that mapping stimuli from fMRI recordings is a significant challenge. This difficulty primarily comes from the complexity of brain activity in the visual cortex, rather than just the volume of data involved [12–14, 20, 23]. Given these insights, it is clear that simple decoding models may not provide accurate reconstructions. To faithfully reconstruct visual images from fMRI data, it is essential to consider and include noise suppression.

To improve model reconstruction and provide better neural explanations, we aim to address the noise problem in linear mapping, leveraging recent advancements in image denoising to guide our efforts. These denoising models can be broadly categorized into three distinct groups. First, we have *Fully-Supervised Denoising Models*, exemplified by deep convolutional neural networks like DnCNN [24], FFDNet [25], CBDNet [26], and SGNet [27]. These models are very effective at removing noise and have been trained using pairs of noisy and clean images. Although they perform well in denoising tasks, they depend on having carefully matched pairs of images for training, which can be difficult to gather in real-world situations. Second, there is the *Noise2Noise* approach, where models are trained with pairs of noisy images. While this method is conceptually interesting, it has fewer practical applications compared to fully supervised methods. Finally, we have *Self-Supervised Models*, a growing approach that is gaining popularity. These models are designed to work with just a single noisy image, as seen in examples like DIP [28] and Self2Self [29]. While promising, self-supervised models also face the challenge of limited applicability. As we explore different denoising models, it becomes clear that each type has its own strengths and limitations. The choice of the best denoising model depends on the specific application and the availability of paired or noisy-noisy data, which will influence future advancements in this field.

In this research, we introduce a new hybrid approach to address the challenges mentioned earlier, combining both linear and nonlinear image decoding methods to reconstruct visual experiences from recorded brain activity. Our initial focus is on linear decoders, which are used to map patterns of brain activity to image pixels. This initial mapping forms the basis for applying nonlinear decoding models, which are then used to denoise and improve the quality of the reconstructed images, resulting in greater clarity and accuracy.

The main goal of this research is to reconstruct visual images from brain activity using both linear and nonlinear decoders, without relying on semantic information. This objective motivates us to investigate the complex relationship between neural activity and visual perception. Additionally, we aim to systematically examine how noise affects the quality and accuracy of our reconstructed images. This dual approach will deepen our understanding of the reconstruction process and shed light on the impact of noise on the results.

## II. Related Work

Visual image reconstruction has recently become a prominent and dynamic area of research, further driven by the significant success of deep learning in computer vision [6, 11–14, 16–20, 23, 30–32]. The effort to accurately reconstruct visual images from neural activity represents a significant intersection of neuroscience and machine learning, presenting a challenging frontier.

This multidisciplinary effort faces several complex challenges from a neuroscientific perspective. Key among these are the large number of voxels and regions of interest, the complexities of encoding and decoding visual information in the brain, and the many aspects of brain function that are still not fully understood [1, 2]. Previous research has shed light on the existence of a meaningful correspondence between visual cortical activities and the perception of visual stimuli [12–14, 16–20, 23]. It has been demonstrated that the experienced visual content can be decoded from fMRI. In the early stages of mapping estimation, linear models, such as linear regression, took precedence. These algorithms typically operate by extracting specific characteristics of the visual content, such as multi-scale local image features, and subsequently learning a linear mapping between these image characteristics and fMRI signals [6]. However, it became clear that linear models faced significant challenges in reconstructing complex images, particularly natural images, as they often produced results with considerable noise.

In contrast, the discovery of similarities between the hierarchical structures of the brain and artificial deep neural networks has led to a new era of effective reconstruction models. Deep neural networks, leveraging this fundamental similarity, have become powerful tools for decoding visual information from brain activity. This shift represents a significant advancement in visual image reconstruction methodologies. For instance, convolutional neural networks (CNNs) are often used to extract features from images and map these features to fMRI signals to reconstruct natural visual content. The development of deep neural network (DNN) architectures has enabled the learning of complex mappings from brain signals to visual content in an end-to-end manner. Various DNN techniques, including Autoencoders, Generative Adversarial Networks (GANs) [33], Recurrent Neural Networks (RNNs) [34], Spiking Neural Networks (SNNs) [35], Diffusion models [32], and Large Visual Language Model (LVLM) [36] have significantly improved the performance of visual image reconstruction from brain activity.

## III. Methods and Results

### A. BOLD5000

Cheng et al. have made the BOLD5000 dataset publicly available [37]. This dataset is a valuable resource for reconstructing visual images from brain activity patterns. It comprises data collected from four subjects (CSI1, CSI2, CSI3, and CSI4), each aged between 24 to 27 years, with either normal vision or vision corrected to normal. These individuals participated in 15 scanning sessions, during which they were exposed to a total of 5254 images. These images include the SUN dataset, consisting of 1000 scene images^3^, the COCO dataset^4^, and the ImageNet dataset, encompassing 1916 images^5^. Combining these diverse datasets enhanced the variety of visual stimuli presented to subjects during scanning sessions. The availability of this dataset and the careful selection of stimuli from multiple sources make it a significant resource for research in visual image reconstruction, as shown in Fig.1.

**Fig. 1:**
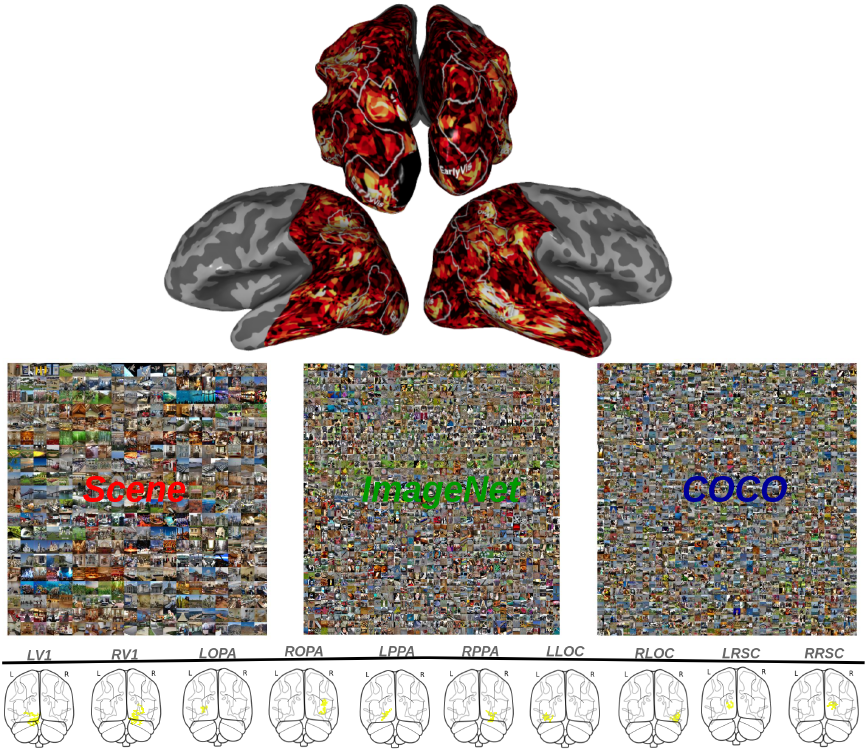
Region of Interest for Natural Scenes. Regions of interest (ROIs) exhibiting the highest color intensity signify specialized regions within the human visual cortex that are selective for scenes, where L refers to the left hemisphere and R refers to the right hemisphere. The brain activity is depicted on a flattened surface, with the corresponding natural scene images positioned in the center.

### B. Functional Localizer Analysis

Participants CSI1, CSI2, and CSI3 underwent eight functional localizer runs across 15 sessions, with up to one localizer run per session. CSI4 had six runs across 9 sessions. The localizer data were analyzed using a general linear model with a canonical hemodynamic response function in SPM12 (https://www.fil.ion.ucl.ac.uk/spm/software/spm12/), including 9 nuisance regressors, regressors for each run, and a high-pass filter. The model included three conditions: scenes, objects, and scrambled images.

### C. Regions of Interest Analyses

Regions of Interest (ROIs) analyses were performed individually using the MarsBaR toolbox with SPM12 in native space. Scene-selective ROIs (parahippocampal place area, PPA; retrosplenial complex, RSC; and occipital place area, OPA) were defined by contrasting scenes with objects and scrambled images. An object-selective ROI (lateral occipital complex, LOC) was defined by contrasting objects with scrambled images. An early visual (EV) ROI was identified by comparing scrambled images to baseline, focusing on the cluster within the calcarine sulcus. ROIs were defined with a threshold of family-wise error correction (p < 0.0001). If activity clusters were too large, a reduced number of localizer runs were used. For scene-selective and object-selective regions, only the first, middle, and last runs were included, while the EV ROI was defined using only the first run due to high activity levels.

### D. Linear and Nonlinear Decoders

#### 1) Linear decoders

Here, we first split all fMRI data and related stimuli into training and validation sets for each subject, and then we considered linear regression [38] as a linear image decoder. Mathematically, we assumed that a linear mapping exists between the latent vectors of visual image stimuli *S* and the corresponding brain response vectors *R*, such that:

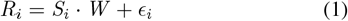

Where *R*_*i*_ ∼ 𝒩 (*β*^*T*^ *R*, *σ*^2^), *ϵ*_*i*_ ∼ 𝒩 (*β*^*T*^ *R*, *σ*^2^), and *i* ∈ ℝ^+∞^ indicate number of stimuli. Training the brain decoder consists of finding the optimal mapping *W* by solving for *W* :

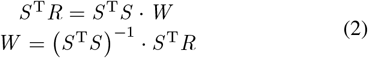

where *S*^*T*^ *S* is the covariance matrix of the latent vectors used for training. Mathematically, testing the brain decoder means using the learned weights *W* to get the latent vector *S* for each new brain activation pattern *R*. Here the weight estimation is performed using *L*2-regularized linear least-squares regression. The optimal regularization parameter for each model is determined by 10-fold cross validation of training data and shared across all voxels. Starting again from Eq.1, we now solve for *S*:

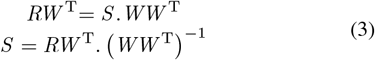

In Fig.2, each linear model predicted voxel responses to new image inputs. This is followed by the integration of the predicted voxel responses from individual models using a weighted linear average. Based on Pearson’s correlation coefficient (Pearson’s correlation between measured and predicted voxel responses) in model training, the weight is calculated for each voxel in the dataset. Overall, we found that linear decoding accuracy varies between individuals, with RSC and OPA having better decoding accuracy than other regions of the high-level visual cortex.

**Fig. 2:**
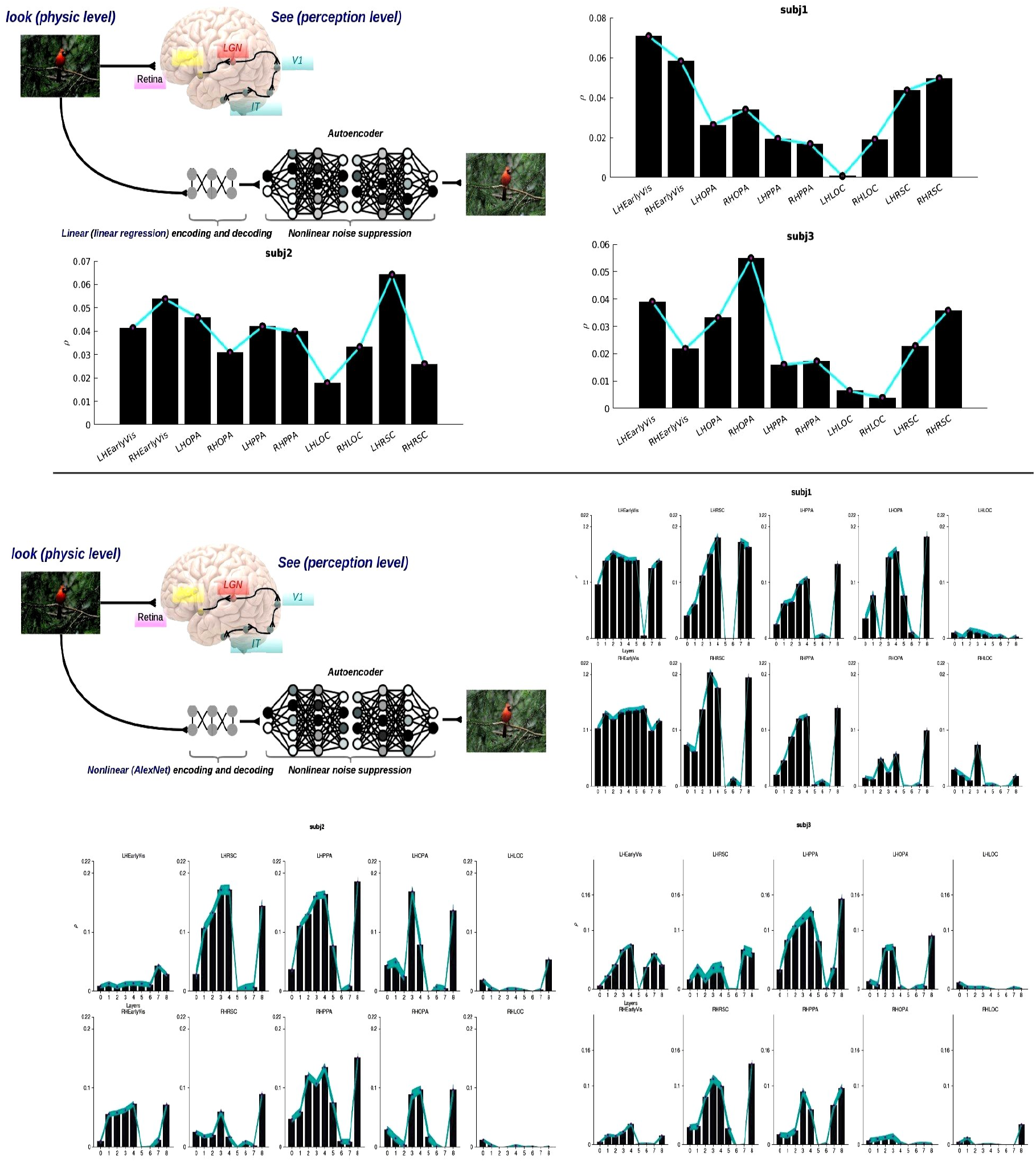
The architecture of linear and nonlinear image decoders, along with nonlinear noise suppression. The top figure provides an overview of the fundamental structure of linear image decoders (linear regression) combined with a nonlinear noise suppression model (autoencoder). Subsequent figures display the linear decoding accuracy observed across various subjects. On the x-axis, we have the regions of interest (ROIs), while the y-axis represents the correlation (*ρ*) between the predicted and actual BOLD responses. These visualizations offer insights into the performance of linear encoding across different ROIs and subjects, with the correlation values (*ρ*) serving as a metric for assessing prediction accuracy in capturing real BOLD responses. In the figure below, the linear image decoder is replaced with the nonlinear image decoder model AlexNet, and the decoding accuracy across different layers is measured for various ROIs and subjects.

#### 2) Nonlinear decoders

The nonlinear deep net model was used to extract the visual image features from each layer of AlexNet [39], more specifically, after the training process, a straightforward feedforward pass involving an input image was used to extract features and perform image denoising using the deep nets. Each unit within the deep nets became active when a natural image was input into the system. The fluctuating representation of a specific feature within the image was captured by an activation weight generated as the image passed through each unit. These units, sharing the same kernel within a single layer, collectively produced an output feature map for each image. The output of each layer was referred to as the result of the rectified linear function before undergoing max-pooling. Then linear regression was applied between the extracted features and brain signals. The related predicted neural responses from different visual regions across different subjects with each layer of deep net were present at the bottom of Fig.2. In general, we see that RSC and OPA high-order visual regions contribute to high decoding accuracy, which totally makes sense considering RSC and OPA are mainly sensitivity-selected responses to complex natural scenes, and the predictive neural response gradually increases with layers from early to middle in deep nets.

#### 3) Color and texture in the visual brain and deep nets

To ascertain the internal features within deep neural networks, we employed the DeepDream methodology as a conventional means for feature visualization in neural networks [40, 41]. This entails forwarding an initialization image through the network and subsequently calculating the gradient of the image with respect to the activations of a designated layer. Subsequently, the image undergoes iterative modifications to enhance the strength of these activations in the context of AlexNet and Autoencoder, employing the Matlab implementation of the DeepDream method^6^. This process serves to depict images that optimally stimulate specific neurons or layers within these networks. These visual representations exhibit a degree of consistency with research asserting functional parallels between the visual cortex and deep neural networks, such as the emergence of progressively more abstract features [42–44].

The Fig.3 reveals that task-demand convolutional neural networks exhibit distinctive features in both low-level and high-level phases of neural networks. For instance, the alexnet and autoencoders primarily manifest white-black (WB), red-green (RG), and yellow-blue (YB) within the convolutional layer 1 (conv1). This pattern aligns with the chromatic-opponent mechanisms found in the primate visual system [45–48] (refer to Figs.3**A**, 3**B**, 3**C**). In layers conv2 and conv3, they feature blob-like texture information while retaining color information, although the chromatic-opponent information gradually diminishes, leading to a more uniform color distribution (Fig.3**A**). In contrast, within the decoder block, all layers predominantly exhibit both color and texture information to effectively reconstruct denoised images. This observation suggests that both color and texture features play pivotal roles in composing images. In essence, color introduces significant changes in the encoder layers, while both color and texture contribute significantly in the decoder blocks to restore clean images. Comparatively, Alexnet networks primarily share color information but diverge significantly in terms of texture information, reflecting their distinct objectives in neural network architecture.

**Fig. 3:**
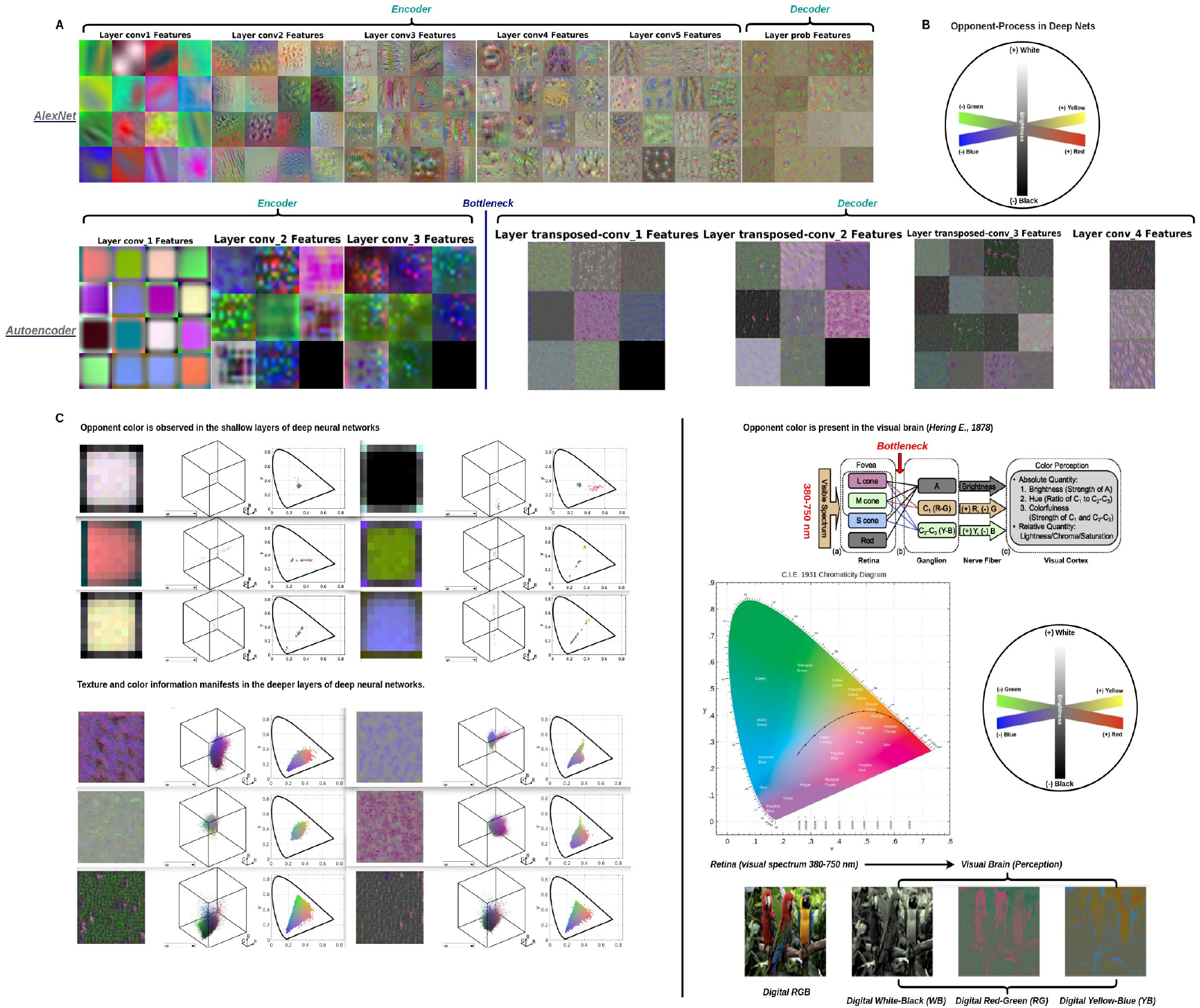
Color and texture information in the visual brain and deep nets. As shown in **A**, the features in AlexNet and the Autoencoder were present in both the encoding and decoding phases. It is clear that color and texture information were captured by both networks. Specifically, color information was primarily retained in the shallow layers of both networks, while texture information was mainly preserved in the deeper layers of AlexNet to enhance image classification. In contrast, the Autoencoder retained both texture and color information in its layers to improve the reconstruction of the original images. The opponent color process is illustrated in **B**, and it is primarily constructed through anti-inhibition pairwise color channels, including white-black (WB), red-green (RG), and yellow-blue (YB). The chosen opponent color distributions in the shallower layers of deep networks are thoughtfully represented in our plots. Specifically, distributions for white-black (WB), red-green (RG), and yellow-blue (YB) are shown in the left panel of **C**. These choices align with the chromatic-opponent mechanisms observed in the primate visual system [47, 48], thereby enhancing the relevance and significance of our visualizations in the right panel of **C**. This panel provides insight into the color distribution of receptive fields in deeper layers, where chromatic-opponent characteristics appear to have diminished, leading to a more uniform color and texture distribution. The figure includes a receptive field visualization, showcasing the intricate interplay of colors within a three-dimensional RGB color gamut. Additionally, a chromaticity diagram is provided to offer a comprehensive understanding of color dynamics, arranged from left to right. Moreover, we present opponent color processing using digital RGB images shown on the right side of **C** to better illustrate the opponent color process in the visual brain [47].

### E. Nonlinear Noise Suppression

In this context, we employed two types of denoised neural networks for our analysis: one set of autoencoder-denoised neural networks that we trained in-house, and another set consisting of a state-of-the-art deep denoised neural network that had already undergone training.

#### 1) Autoencoder-denoised neural networks

In this study, we harnessed the power of denoised Convolutional Neural Networks (CNNs), a cutting-edge advancement in artificial intelligence known as Autoencoder [50]. These networks were utilized to address denoising challenges, particularly for images degraded by Gaussian white noise. The core concept behind Autoencoders lies in having the generator’s target coincide with its input. Essentially, the generator’s mission is to encode the input data into a compressed latent representation and then proficiently decode it back into the original images [50]. Furthermore, it shares both structural and functional similarities with the human visual system [31, 49]. Furthermore, the previous study shows that visual brain performance enhances image quality processing when input images are of low quality [49], as shown in Fig.4**A**.

**Fig. 4:**
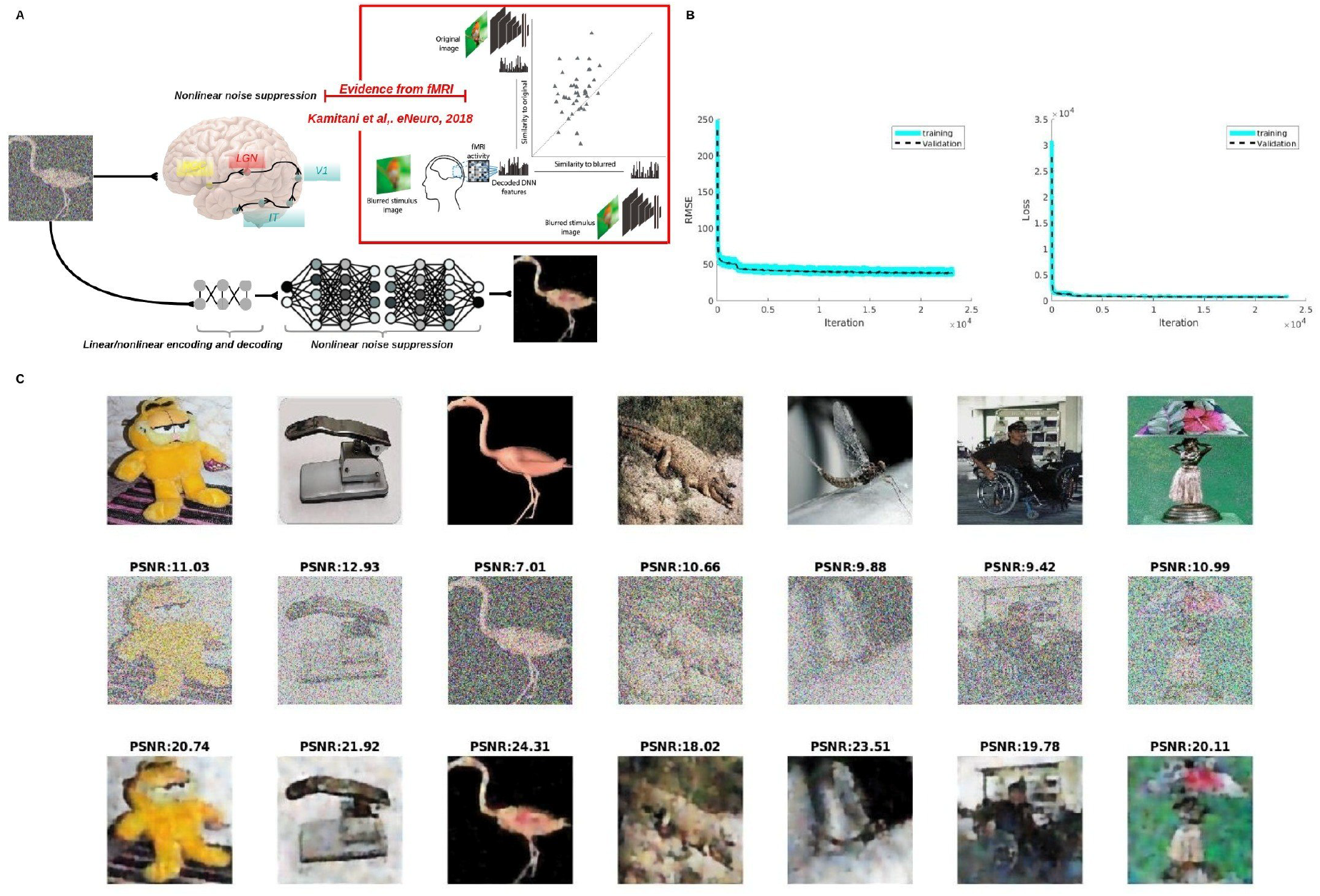
Nonlinear noise suppression in the visual brain and deep nets. Previous fMRI experiments have revealed that the visual brain performs denoising processes [49], potentially enhancing images affected by noise. In this study, we trained a deep autoencoder to suppress noise, taking advantage of the structural and functional similarities between the visual cortex and the autoencoder. The training and validation curves are shown in **B**, while the denoising performance, quantified using peak signal-to-noise ratio (PSNR), is illustrated in **C**.

To facilitate the training process, we leveraged the Caltech-101 dataset [51], a comprehensive collection comprising 101 distinct classes, including a wide range of objects such as animals, airplanes, and flowers, among others, each exhibiting high shape variability. Notably, the dataset encompasses varying numbers of images per category, ranging from 31 to 800, with most images characterized by medium resolution, approximately 300×300 pixels. In preparation for network training, we resized the input images to dimensions of 224 by 224 pixels.

To introduce controlled noise and further enhance the diversity of our training and validation samples, we intentionally added Gaussian white noise to create degraded images. Specifically, we applied noise with a mean value of 0.5 and a variance of 0.6. Additionally, we employed data augmentation techniques to augment our training and validation datasets, enriching the learning process and enhancing the network’s ability to handle noise and variations effectively.

After that, we trained networks on images using a regression output layer architecture with a six-layer encoder plus a six-layer decoder (convolution layer with 16 filters of size [3 3], max pooling layer with pool size [2 2] and stride [2 2], convolution layer with 8 filters of size [3 3], max pooling layer with pool size [2 2] and stride [2 2], convolution layer with 8 filters of size [3 3], max pooling layer with pool size [2 2] and stride [2 2]), and the decoder component of the architecture mirrors the encoder (upsample transposed convolution layer with 8 filters of size [2 2], upsample transposed convolution layer with 8 filters of size [2 2], upsample transposed convolution layer with 16 filters of size [2 2], convolution layer with 3 filters of size [3 3]). We trained neural networks with their loss function to optimize mean-squared-error (MSE). All layers are convolutional or up-convolutional. Because the autoencoder networks work with color images and accept input images of fixed sizes, if the input images were grayscale images, then we converted them into RGB images by stacking three copies of the grayscale image G to create a corresponding 3-D array RGB = [G G G], and resizing them to fit into the input image size of the autoencoder network. The input image size for the autoencoder is 224-by-224. For the training options, the ADAM stochastic gradient descent was used. The momentum value was set to 0.9000, and the gradient threshold method was set to *L*2 norm. The minimum batch size was 800, and the maximum number of epochs was 1, 000. The initial learning rate was 0.001, and it stayed the same throughout training. The training data were shuffled before each training epoch, and the validation data were shuffled before each network validation. The final denoise performance of deep nets is shown in Figs.4**B**,**C**.

#### 2) Pre-trained state-of-the-art deep convolutional denosied neural networks

In our study, we harnessed the power of a pre-trained deep convolutional neural network known as DUBD to effectively remove noise [52]. This choice was intentional, as the DUBD model has consistently outperformed other state-of-the-art denoised neural networks. For a comprehensive understanding of the pre-trained network’s architecture and loss function, we refer readers to the detailed description provided in [52].

Our experimentation involved subjecting images to varying noise levels, specifically 35, 60, 96, and 130, in order to assess the denoising performance of the DUBD model. We quantified the model’s effectiveness in reducing noise and enhancing image quality using the Root Mean Square Error (RMSE) metric, as shown in Fig.5**A**. Meanwhile, we presented the results of visual image reconstruction from various brain regions using linear regression in combination with different levels of nonlinear denoising processes. The findings indicate our capability to predict and reconstruct visual images based on brain activity, devoid of the incorporation of semantic information. Specifically, we have put together a selection of visually impressive reconstructions, average reconstruction images, and reconstructions with notably diminished quality, as shown in Figs.5**B**, 5**C**.

**Fig. 5:**
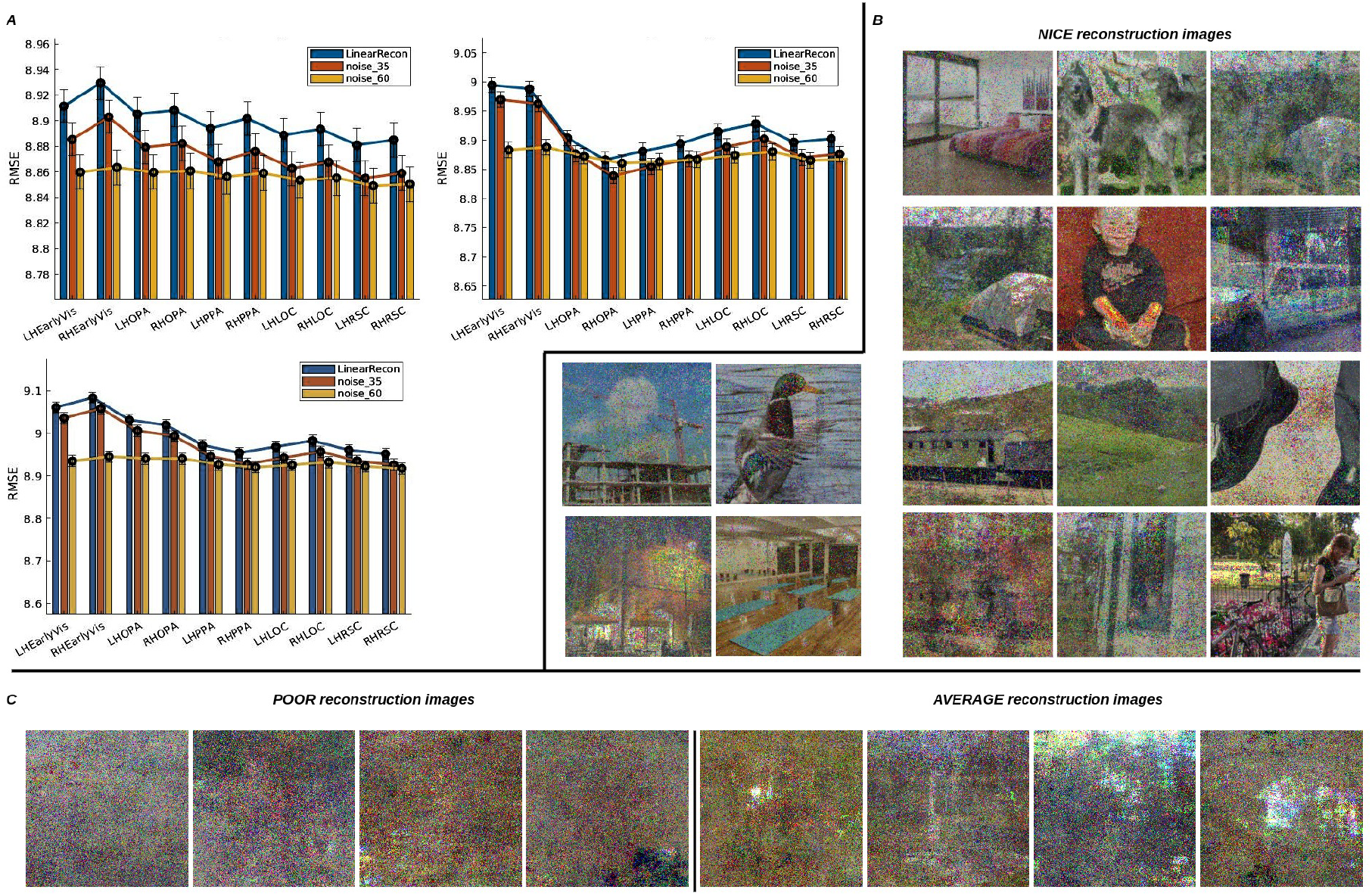
Visual image reconstruction from brain activity. Visual image reconstruction quality was evaluated for various subjects using linear regression combined with nonlinear denoising across different brain regions, as shown in **A**. We report the Root Mean Square Error (RMSE) for various brain regions reconstructed across these subjects. In each mini-group, the first bar represents linear reconstruction without denoising. The second bar shows denoising using dCNN with a noise level of 35 applied after linear reconstruction. The third bar reflects the same processing with a noise level of 60, which is the optimal choice for suppressing noise. These data provides valuable insights into the impact of denoising on image quality across different brain regions. High-quality reconstruction images from brain activity, selected across different subjects, are shown in **B**. Poor and average reconstruction images from brain activity, selected across different subjects, are shown in **C**.

To further highlight the superiority of our models, we compared our reconstruction performance with that of Variational Auto-Encoder (VAE/GAN) and Wasserstein Auto-Encoder/Dual-Variational Autoencoder (WAE/Dual-GAN) [15]. As shown in Fig.6, it is clear that our reconstruction results exhibit the best performance compared to those of GAN reconstruction.

**Fig. 6:**
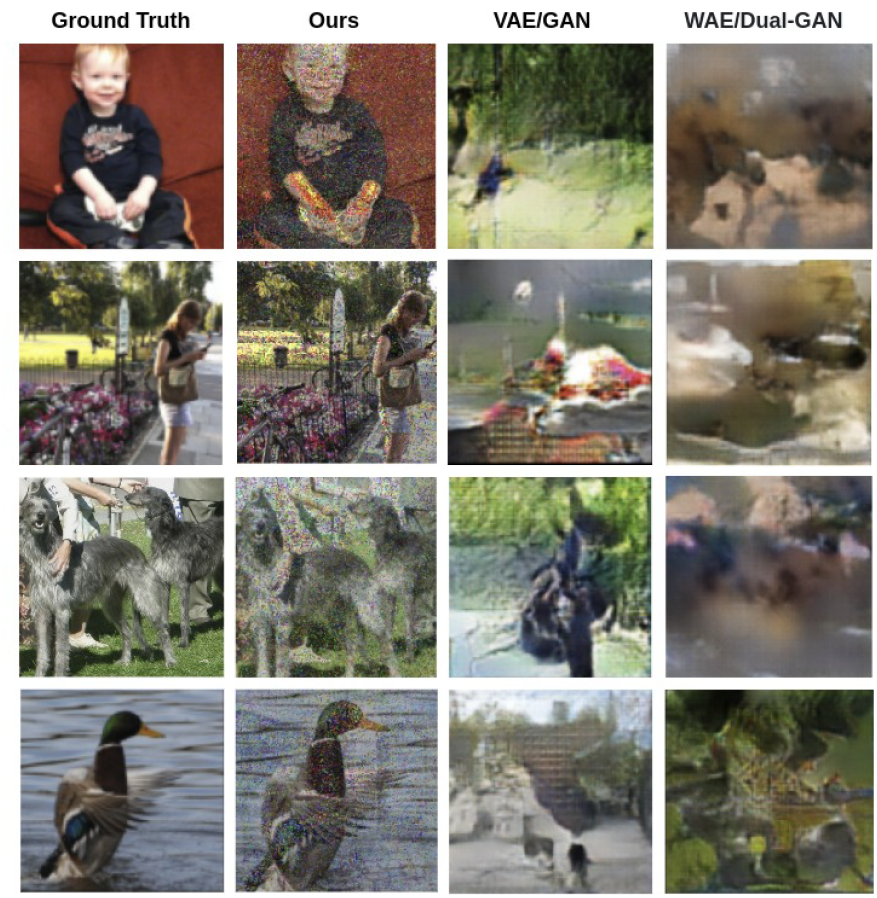
Reconstruction performance using different methods. The first column shows the ground truth images, followed by our reconstruction images, and then the reconstruction results from VAE/GAN and WAE/Dual-GAN.

## IV. Conclusion and Discussion

Our findings demonstrate that noise can be effectively reduced and reconstructed image quality from brain activity enhanced when using linear regression in conjunction with a denoising process. Thus, it is possible to reconstruct visual images from brain activity without relying on complex diffusion models or text-to-image methods. Although text-to-image methods can achieve impressive performance, they might overlook the underlying neurological mechanisms, as we do not fully understand the inner workings of these models. Additionally, we observed that the high-level visual cortex showed fewer errors in image reconstruction compared to the low-level visual cortex. Biologically, this observation is consistent with our understanding that natural images, such as objects, scenes, and complex visual content, which are typically represented in high-level brain regions.

Furthermore, we examined the inner workings of the deep networks used in our proposed model and observed that some low-level properties, such as color and texture representation, align with those in the visual brain. This alignment provides clearer neural explanations compared to other complex or semantic-guided models. Our model, which aligns both structurally and functionally with the visual brain, not only offers impressive reconstructions but also maintains strong neural explanatory power.

In our future research endeavors, we may explore the application of principal component analysis or salinecy mapping to complex nature images to reduce image redundancy, potentially leading to improved reconstruction performance. Additionally, it’s worth noting that the suboptimal reconstruction results observed in this study could be attributed to the relatively low Signal-to-Noise Ratio (SNR) in the BOLD5000 dataset. Therefore, we should consider utilizing datasets with higher SNR levels, such as the Natural Scene Dataset [53], to potentially enhance our outcomes. Furthermore, it is imperative to investigate the interactions within high-order visual regions during task states, as this exploration has the potential to enhance decoding accuracy. High-order interactions are known to provide richer information compared to low-order brain networks [9, 54, 55]. Simultaneously, we must take into account bidirectional information interactions within the visual brain during visual decoding. This consideration may not only lead to improved decoding accuracy but also facilitate a more profound understanding of the information flow within the visual system during visual tasks [56, 57]. Lastly, we should also investigate the potential of deep approximation neural networks, as they have demonstrated remarkable achievements in various domains, and could be a promising avenue for future research.

## V. Abbreviations

BOLD: Blood-Oxygen-Level-Dependent
fMRI: Functional Magnetic Resonance Imaging
CNNs: Convolutional Neural Networks
DNN: Deep Neural Network
GANs: Generative Adversarial Networks
RNNs: Recurrent Neural Networks
SNNs: Spiking Neural Networks
L: Left Hemisphere
R: Right Hemisphere
EV: Early Visual
PPA: Parahippocampal Place Area
RSC: RetroSplenial Cortex
OPA: Occipital Place Area
LOC: Lateral Occipital Cortex
ROIs: Regions of Interest
VAE: Variational Auto-Encoder
Dual-GAN: Dual-Variational Autoencoder
WAE: Wasserstein Auto-Encoder

## Footnotes

1 Short-term memory, also known as primary or active memory, is the ability to hold a small amount of information for a short period.

2 Long-term memory is the ability to store information for a long time, often for many years.

3 http://sun.cs.princeton.edu/

4 https://cocodataset.org/

5 http://www.image-net.org/

6 https://www.mathworks.com/help/deeplearning/ref/deepdreamimage.html

